# KS-Burden: Assessing distributional differences of rare variants in dichotomous traits

**DOI:** 10.1101/368696

**Authors:** Robert M. Porsch, Timothy Mak, Clara Tang, Pak C. Sham

## Abstract

A number of rare variant tests have been developed to explore the effect of low frequency genetic variations on complex phenotypes. However, an often neglected aspect in these tests is the position of genetic variations. Here we are proposing a way to assess the differences in spatial organization of rare variants by assessing their distributional differences between affected and unaffected subjects. To do so, we have formulated an adaptation of the well know Kolmogorov-Smirnov (KS) test, combining both KS and a simple gene burden approach, called KS-Burden.

The performance of our test was evaluated under a comprehensive simulations framework using real data and various scenarios. Our results show that the KS-Burden test is able to outperform the commonly used SKAT-O test, as well as others, in the presents of clusters of causal variants within a genomic region. Furthermore, our test is able to maintain competitive statistical power in scenarios unfavorable to its original assumptions. Hence, the KS-Burden test is a valuable alternative to existing tests and provides better statistical power in the presents of causal clusters within a gene.

## Introduction

The advent of genome-wide association studies (GWAS) has contributed significantly to our understanding of complex traits by finding several thousand robust association between genetic variants and complex phenotypes [1]. However, GWASs survey only common variants (*MAF* > 0.01) and ignore lower frequency variations which make up the majority of polymorphisms. Detection of low frequency variants have been more challenging but the recent development of next-generation sequencing based technologies have provided rich opportunities to study those rare variants and their impact on complex human traits [2].

Indeed, rare variants, which can be defined as genetic variations occurring in less than 1% of the population, have been suggested to play an important role in the etiology of human traits and potentially account for the missing heritability [3, 4]. Thus considerable effort has been made to develop and deploy statistical methods to discover important causal relationships between rare variants and complex human traits [5–8]. In GWASs, a single variants is associated with the trait in question. This approach is largely unfeasible in rare variants due to their low frequency and large numbers, as well as the limited sample size of most studies [9]. Thus most approaches have been focused on combining multiple rare variants in order to increase statistical power. This can be either done on the gene or pathway level, but for simplicity we will only consider gene based tests within this paper.

In general, one can classify rare variant tests into three categories, namely burden, variance-component and omnibus tests, based on their assumption regarding the underlying genetic architecture [9]. In general, burden tests aggregate single rare genetic variations. Thus assuming that all variants in a given genomic region have the same direction of effect. Violation of this assumption results in considerable reduction in statistical power. Examples of this methods include the Combined Multivariate and Collapsing (CMC) test [10], as well as the weighted sum statistic [11]. Alternatively, variance-component tests do not assume uni-directional effect of all included variants. These methods investigate the distribution of genetic effects for a genomic region and are robust to variants with differing direction of effects. Prominent example of variance-component tests are SKAT [12] and C-Alpha [13]. These tests are more powerful compared to the burden approach in situations when the majority of rare variants are of neutral or bi-directional effect. However, burden tests in general outperform variance-component based tests when a large proportion of variants have the same direction of effect. This has lead to the development of omnibus tests to combine both approaches. A commonly used example of these omnibus tests is SKAT-O [14] which uses a combination of SKAT and burden tests statistics to derive a combined p-value.

An often neglected aspect of rare variant tests are the position of these genetic variation and only a few tests have so far been suggested [15–17]. Multiple biological evidence has been reported in the past demonstrating clustering of causal rare variants within the genome [16]. It is biological plausible to suggest that rare deleterious mutations causally related to a considered trait might be more likely to be located in protein functional domains or gene regulatory elements.

We are here proposing a way to assess the differences in spatial organization by assessing the distributional differences of rare variants between cases and controls. To do so we make use of the well known Kolmogorov-Smirnov (KS) test. We demonstrate, through simulations, that our methods shows good statistical power compared to commonly used tests, such as SKAT and burden, when the assumption of the KS test are met. Further, we combine the KS and burden test to provide an omnibus approach to our gene based association tests.

## Materials and methods

### Notation

Assume *n* subjects are sequenced (indexed by *i*) and in a given region and *p* variant sites were observed (indexed by *j*). Then *G_i_* = (*G_i_*_1_*, …, G_ip_*) denotes the genotype for the *p* variants. One assumes an additive genetic model and *G_ij_* = 0, 1, 2 represents the number of copies of the minor allele.

### Kolmogorov-Smirnov (KS) Test

The two sample KS test is a non-parametric test for the equality of two one-dimensional probability distributions. The test can be adapted to test for the distributional differences of multiple variants in a region on a dichotomous phenotype by computing the respective cumulative distribution functions (see Figure 1).

**Fig 1.**
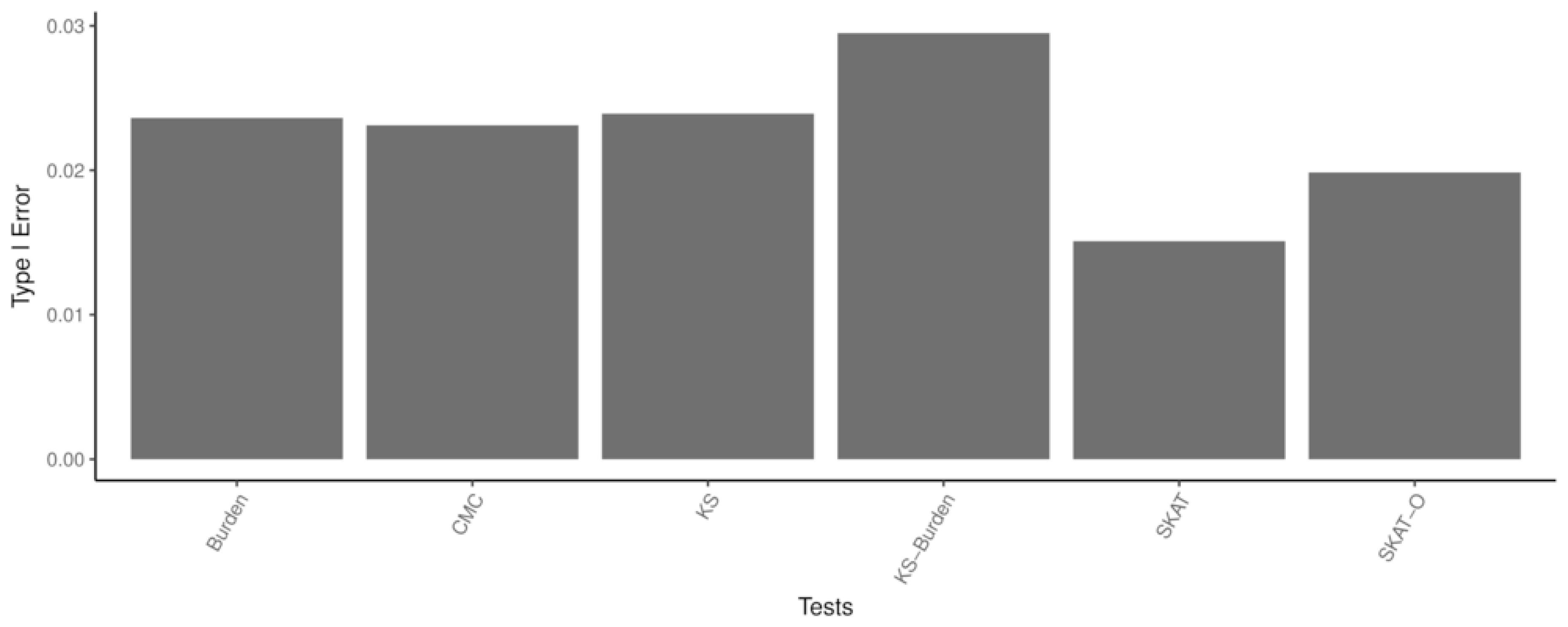
Illustrative example of the KS test applied to a genomic region. The arrow indicates the larges distance between the two cumulative distribution functions of affected and unaffected individuals.

For a given genotype matrix *G* of size *n* × *p* one can compute the empirical cumulative distribution function at any given variant position *x* in *G* as

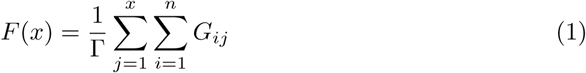

in which 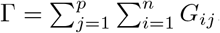. Hence *F* (*x*) computes the cumulative proportion of alleles present in a region from the beginning up until the position *x* within a genomic region.

Given the genotype matrices of affected and unaffected individuals (*G_A_* and *G_U_*) on can compute *F* (*x*) for both groups separately for all genomic positions *x*. The test statistic of the KS test is then given as

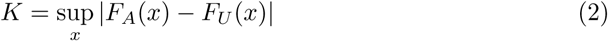

Thus the KS test aims to identify distributional differences of multiple variants by identifying the largest absolute differences between the two empirical cumulative distribution functions.

Evaluating distributional differences of variants between affected and unaffected individuals corresponds to testing the null hypothesis *H*_0_ : *F_A_* = *F_U_*. While the classical KS test is distribution-free when *F* is continuous, it is not the case with discrete data, such as allele count. Indeed, application of the Kolmogorov distribution to obtain critical values for *K* for discrete data yields conservative estimates [18, 19]. Therefore, we applied a permutation-based approach to evaluate *K* under *H*_0_.

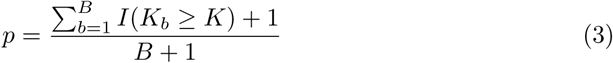

In which *K_b_* is the test statistic of the *b^th^* permuted sample and *K* is the observed test statistic.

### Omnibus Test for KS and Burden

The KS test evaluates distributional differences of multiple variants in a given region, but does not test for overall differences in the amount of variants between affected and unaffected individuals. The Burden test, in contrast, does not take distributional differences into account but tests for the differences in allele counts between affected and unaffected individuals. One can define the test statistic of the Burden tests as

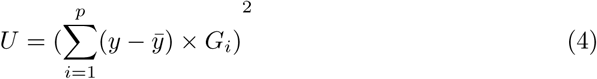

Hence, under *H*_0_ the KS test and Burden test can be seen as orthogonal to each other. Multiple different methods have been developed to combine two different tests, most prominently Fisher’s product methods. We made use of a similar developed which has been shown to be more powerful [20]. Specially, given the ordered p-values of both KS and Burden (indexed by *j*) one can compute *W_j_* as

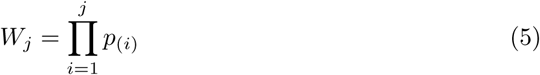

Hence, *W*_1_ represents the smallest p-value from either KS or Burden, while *W*_2_ is the product of both p-values.

This combined test rejects the null if 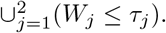 The critical value *τ_j_* for *W*_1_ and *W*_2_ is estimated via a Monte Carlo approach as following:

1. Create a matrix *P* sized *L ×* 2 with elements *p_lj_, j* = 1, 2; *l* = 1,…, *L* for some large integer *L*, in which *p_lj_* ~ *U* (0,1)
2. Calculate *p_l_*_(1)_ and 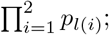; denote them as *W*_[*l.*]_ = *w_l_*_1_*, w_l_*_2_
3. Compute the *α*-percentile for each column of *W* and note the critical values *τ_j_* This relatively simple approach allows to combine both KS and Burden, while holding the type 1 error rate stable [20].

Implementation of the KS, Burden and the combined KS-Burden was done in C++ and can be found at https://github.com/rmporsch/ksburden.

## Simulation Study

In contrast to most previous studies investigating the statistical properties of rare variant association tests, we made use of real sequencing data. Thus, genotypes were not simulated but acquired from a previous sequencing study. The main aim of using real data is to accurately reflect the diversity of genes within the human genome as well as limitations commonly encountered in sequencing-based studies. Indeed, most gene based tests are confronted with a variety of small and medium gene lengths which is often not reflected in the original power estimations. Hence we will make use of a large sequenced sample originally intended to study Hirschsprung’s diseases to investigate the statistical properties of the KS and KS-Burden test.

### Simulation Dataset: The Hirschsprung Sequencing Project

Hirschsprung disease (HSCR) is a rare congenital disorder occurring in about one in 5, 000 births [21]. The here used sample contains 464 cases as well as 498 controls analyzed by whole-genome sequencing (36.6X median read depth). All patients are sporadic cases with known family history of HSCR and were recruited in China (*n* = 341) and Hanoi, Vietnam (*n* = 102). Controls were obtained from same or nearby cities in order to match the subpopulation of patients. Samples were excluded due to failure in heterozygosity (an excessive amount of heterozygous genotypes in a sample can indicate potential genotyping problems), gender accordance, duplications, and relatedness. Population outliers, e.i. samples which were not of Chinese or Vietnamese descent, were removed by PCA resulting in a final set of 443 cases and 493 controls. Furthermore, genotype level quality control was applied with KGGseq [22] and genotypes with quality less than 20 (*GQ* < 20) or covered by less than 8 reads (*DP* < 8) were excluded. In addition, variants with call rate < 0.9 or those violating Hardy-Weinberg equilibrium (*p <* 10^−5^) were removed. This resulted in a final call set of 33.5 million (M) SNVs and 3.3M indels, the majority of which are novel (61.5% for SNVs and 68.7% for indels).

Variants were then annotated for protein functions against RefSeq gene annotations as well as population frequencies (1000 Genomes Project phase 3, ExAC and ESP). Only nonsynonymous exonic variants (frameshift, nonframeshift, stopgain, missense, startloss, stoploss, splicing) with an MAF ≤ 1% were included in the simulations. Overall frequency of rare nonsynonymous mutations per gene are displayed in Figure 2.

**Fig 2.**
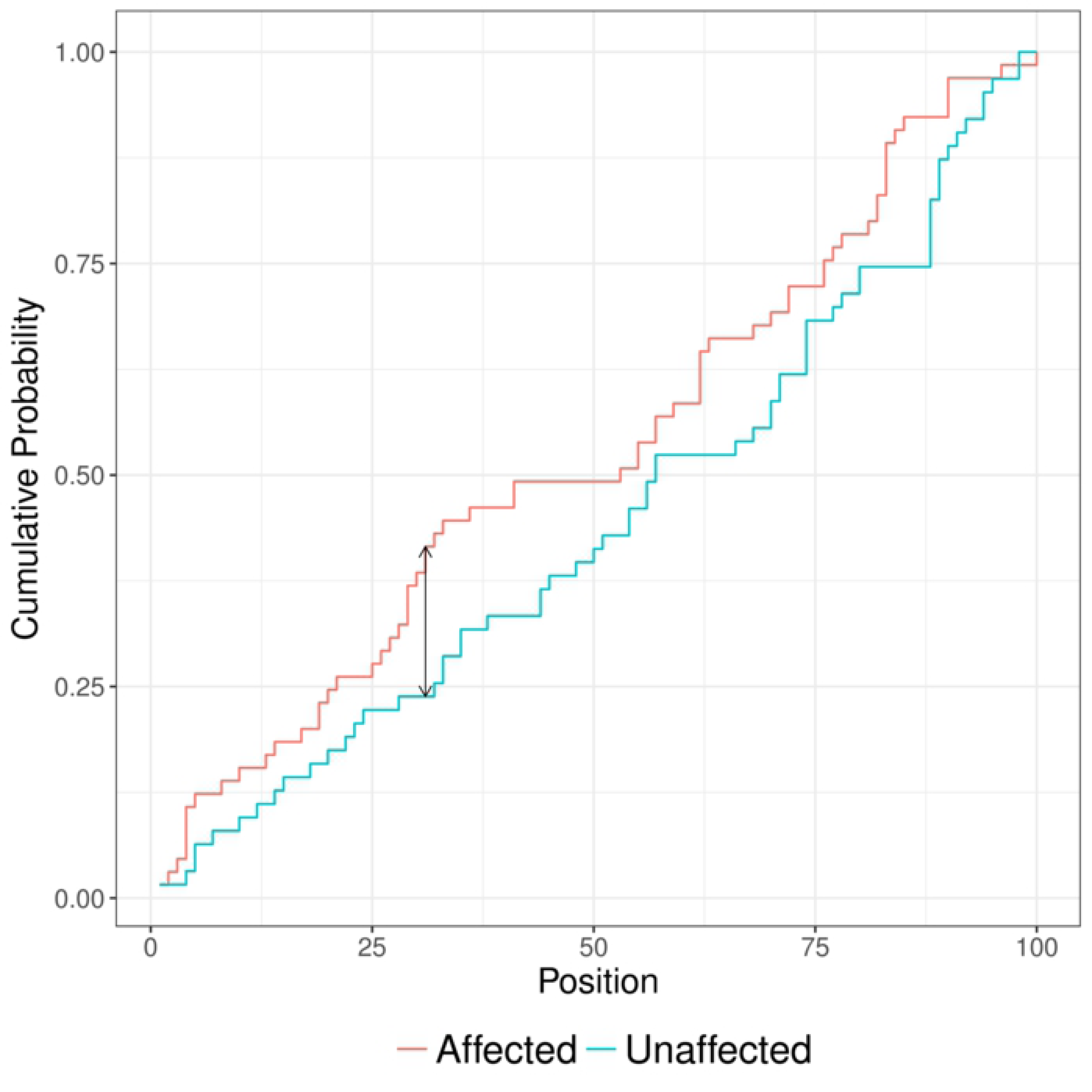
Overall frequency of rare non-synonymous mutations per gene in HSCR data set.

### Simulation Framework

To explore and test the statistical properties of the KS and KS-Burden test we carried out simulations under a number of different scenarios. For all simulations we used sequenced genotypes from our seed population sample. From the set of 15, 480 genes present within this population we selected 50 genes at random which had at least 10 or more rare variants. Variants were defined as rare if the minor allele frequency (MAF) was below or equal 1%. We simulated two different scenarios (see Figure 3 and Figure 4) in which we assume different configurations of causal clusters *γ*.

**Fig 3.**
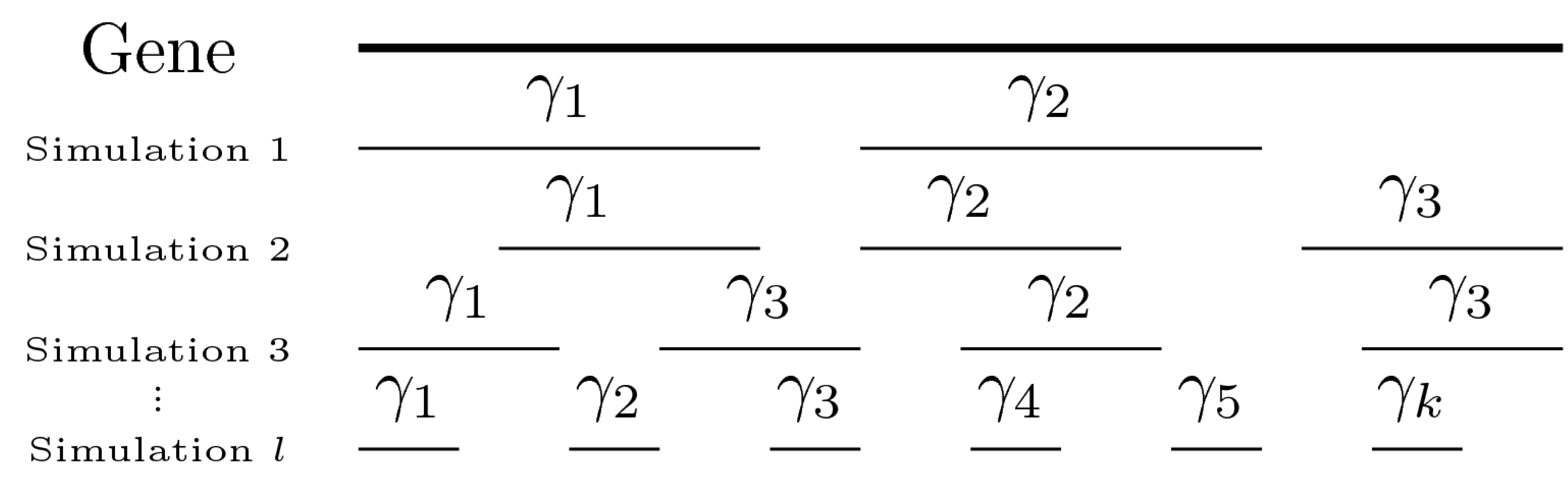
Graphical representation of the simulation configurations 1

**Fig 4.**
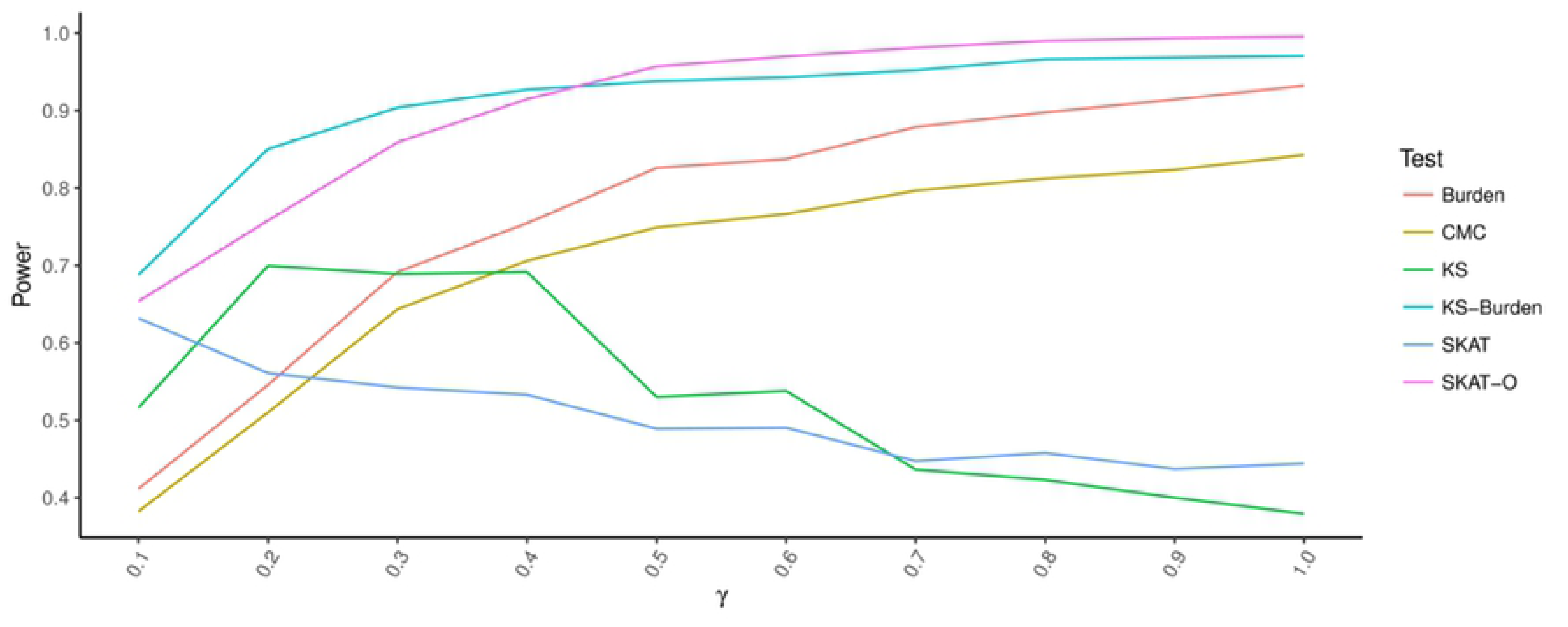
Graphical representation of the simulation configurations 2

The phenotype for each configuration was simulated via a liability threshold model. Hence the phenotype *Y_i_* of the *i^th^* subject was generated via 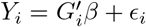 in which *G_i_* are the standardized genotype of the nth subjects with *P* variants, *β* is the effect size vector of size 1 × *P* and *ϵ _i_* is a standard normally-distributed error term with a mean of 0 and a variance of 1 – *h* in which 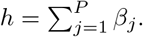 The effect h was uniformly distributed across all causal variants and therefore representing the effect of the whole genomic region. We assigned case status for each subject whose *Y_i_* was above a certain liability threshold, *q*. This process was repeated until 500 cases and an equal number of controls were collected.

### Configuration 1

We assumed a single cluster in a given genomic region, called *γ*, which is located at random positions within the genomic region. All variants in *γ* were assigned to be causal. Furthermore, the size of *γ* was expressed as the proportion of variants included in the causal cluster relative to the total number of variants in a given region. For example, given *γ* = 0.1 and a gene with 100 rare variants causal status would be assigned to a cluster of 10 variants.

### Configuration 2

In addition to randomly placing a single causal cluster within the genomic region, we extended the number of clusters to more than 1. Variants in all clusters were assigned to be casual, while the proportion of causal variants within the genomic regions (*γ*) were kept constant.

## Results

For each gene and scenario, statistical power was evaluated after 1000 replications, with a liability threshold *q* of 0.01 and an effect size *h* = 0.02. The significant threshold *α* was set to 0.05. Overall statistical power was then estimated as the average power across 50 randomly chosen genes.

As expected, both KS and KS-Burden performed favorably, compared to other commonly used gene based tests in the presents of a single causal cluster. Specially, the KS test shows stable statistical power at *γ* = {0.2–0.5}, while declining at larger causal cluster sizes. Interestingly, at *γ* = 0.1 power of the KS test was reduced, possible due to the limited length of some genes. The omnibus KS-Burden test shows superior performance between *γ* = 0.1 and *γ* = 0.5 compared to SKAT-O as well as others, while maintaining similar statistical power at larger cluster sizes. The two burden tests, CMC and Burden, perform poorly in the presents of small clusters, but retain competitive power at larger cluster sizes. Interestingly, the simple Burden test seems to underperfrom relative to SKAT-O in scanarios most favorable to the former. Specially, the Burden test assumes that all variants have the same direction of effect, a situation which corresponds to *γ* = 1.0. While within this scenario the Burden test does indeed expresses its highest statiscal power, SKAT-O as well as KS-Burden are able to outperform the simple Burden test.

To conclude, given a single causal region, the KS-Burden test is a valuable alternative to the commonly used SKAT-O. Furthermore, out test demonstrates better statistical power when a single small cluster of causal variants is present, while maintaining good performance at larger cluster sizes.

Next, we explored the behavior of the our developed tests in situations with more than 1 causal cluster (see Figure 5). Not surprisingly, statistical power of the KS test is reduced with the increase of causal clusters while holding the total size of clusters relative to the size of the genomic region constant. In contrast KS-Burden is not affected by the increase in the number of clusters. Similar our simulations show that the statistical power of CMC, Burden, SKAT and SKAT-O is independent of the number of clusters.

**Fig 5.**
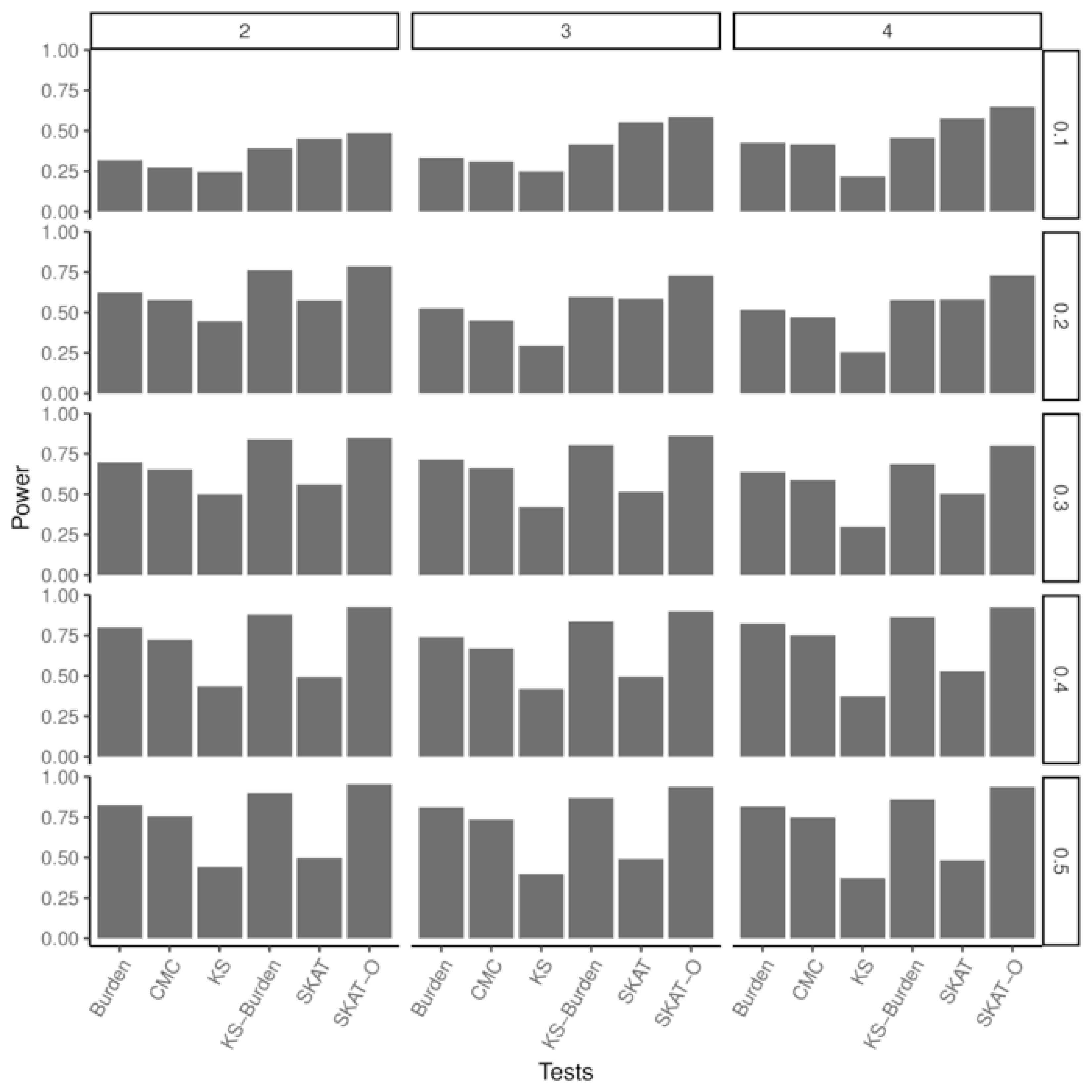
Estimated statistical power for four different causal cluster size *γ*.

**Fig 6.**
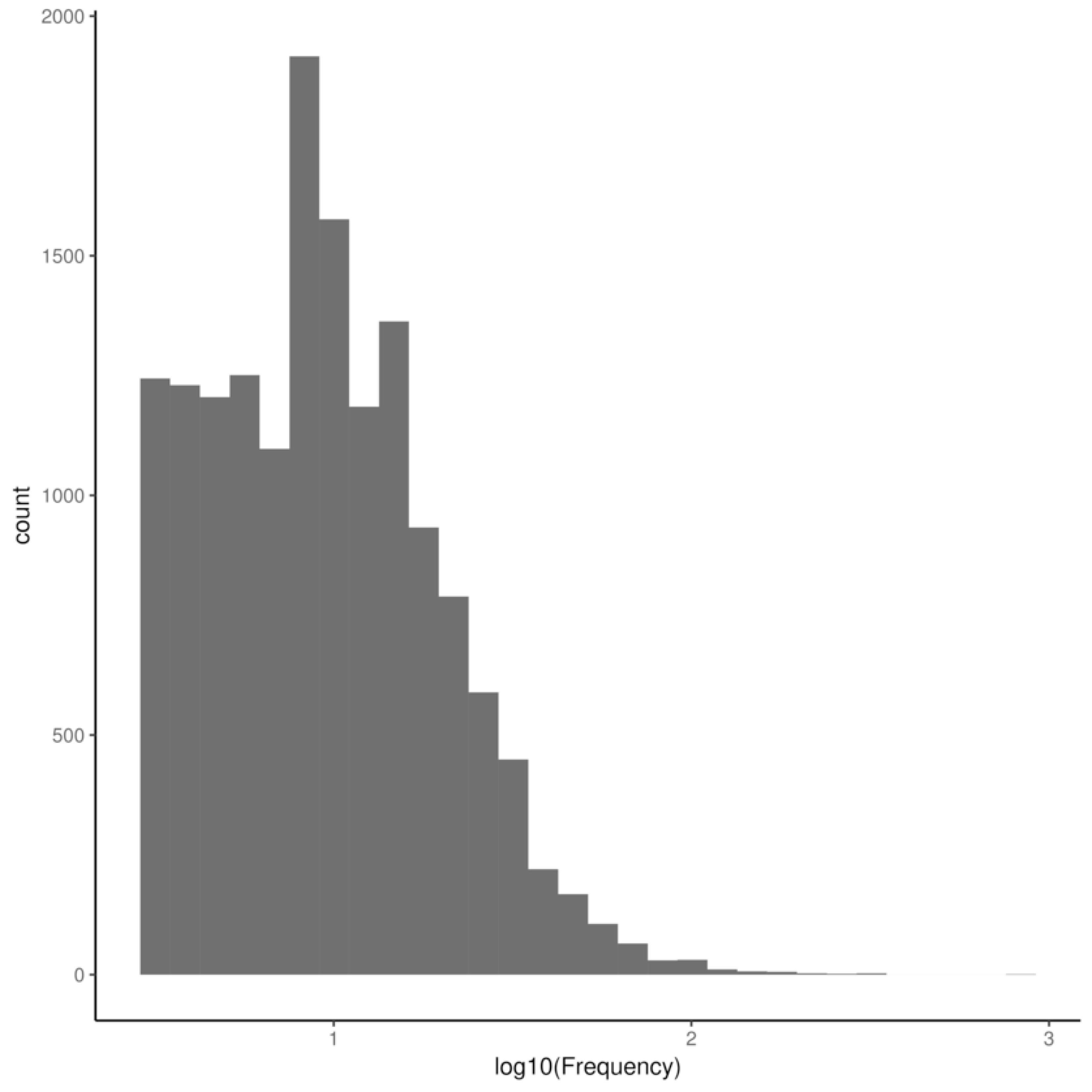
Statistical power for Burden, CMC, KS, KS-Burden, SKAT and SKAT-O for different cluster sizes (right panel) and number of clusters (top panel).

Overall, SKAT-O performs favorable in most scenarios, while KS and KS-Burden lose statistical power relative to the other tests as the number of causal clusters increases. It is notably, however, that the KS-Burden test is able to maintain good power in many scenarios unfavorable to the KS test. Demonstrating that our omnibus approach is able to compensate for the shortcomings of the KS test.

Hence, these simulations have shown that the KS-Burden is able to compensate mismatched assumptions of the KS test effectively and that the KS-Burden is able to maintain appropriate performance across a variety of scenarios.

Overall, our simulations have shown that the KS-Burden test is able to outperform commonly used tests in some specific scenarios. Furthermore, the test is able to maintain good statistical power in simulations unfavorable to the KS test. Therefore, providing a valuable alternative to commonly used tests.

Interestingly, type 1 error rate for all tests is significantly lower than the chosen *α* of 0.05 (see Figure 7). It is important to note that permutation approaches are known to estimate conservative p-value in rare event data, such as rare genetic variations. Nevertheless, type 1 error rate is noticeable low for both SKAT and SKAT-O, while the burden based approaches, such as CMC and Burden are slightly higher. The highest type 1 error rate is present in our KS-Burden test, but still significantly below 0.05.

**Fig 7.**
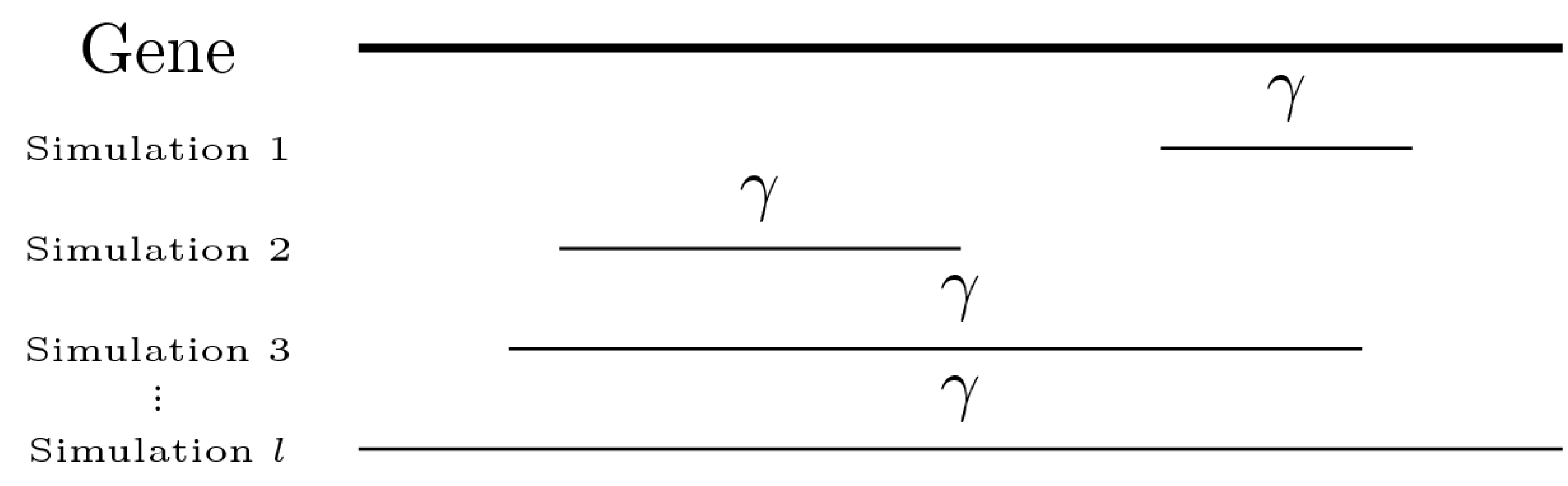
Type 1 error rate for various rare variant tests.

## Discussion

We have shown that the KS-Burden test is able to outperform commonly used gene-based tests when a single causal cluster is present within a genomic region. Furthermore, our test shows similar performance compared to other tests in scenarios which are unfavorable to its underlying assumptions.

It is important to emphasize all gene based tests make assumptions about the underlying genetic architecture [9]. Specifically, while the Burden tests assumes that all variants have the same direction of effect and SKAT assumes the presents of protective and damaging alleles, the KS test makes the assumption that a single causal region is present in the genomic region under consideration. These underlying assumptions reflect the researchers prior believe about the underlying genetic architecture. For example, given a well known genomic region, which has been exhaustively annotated with information about the potential pathogenicity of all contained variants, one would expect that most, if not all, variants selected for a gene based rare-variant test have the same direction effect. Hence the Burden test would yield the best statistical power in this particular situation. On the other hand, an ill defined genomic region with little to no variant annotations would be more favorable to both KS and SKAT since many variants would be of neutral effect towards the phenotype. Here it is important to note that the KS test makes the reasonable biological assumption that causal variants are clustered together. Thus increasing statistical power in the presents of such cluster. In the absence of such cluster, the KS-Burden test was designed to compensate for such shortcomings, similar how SKAT-O combats assumption mismatch of SKAT. Importantly, the KS-Burden test is able to maintain the good statistical power in the present of assumption mismatch, while still accounting for the potential presents of a single causal cluster. This makes the KS-Burden test a good candidate for rare variant testing in genomic region with little or no biological annotations. Furthermore, given that the KS test statistic is the largest absolute difference between the two empirical cumulative distribution functions the KS test is able to provide researchers with a specific location of the causal cluster. Thus not only providing better statistical power when the assumptions of the test hold, but also delivering valuable biological insight.

In addition, the low type 1 error rate across all tests needs to be discussed which is in contrast to previous studies [12–14, 23, 24]. Interestingly, most previous studies did only used simulated genotype matrices to estimate statistical power. In contrast, our study made use of a large whole genome sequencing data set therefore reflecting commonly encountered scenarios in rare variant association studies. Indeed, most genes used in our analysis are relatively small and contain only a few rare variants. This has a direct effect on the commonly used permutation approaches and results in conservative p-value estimations. However, this issue is present across all used tests and should be reduced in larger sample sizes.

Furthermore, it is somewhat surprising that the Burden test is unable to outperform SKAT-O and KS-Burden in situations most favorable to it (*γ* = 1.0). However, it is important to note that even under very unfavorable scenarios both KS and SKAT are able to retain some statistical power which has not been captured by Burden. Hence explaining the superior performance of the two omnibus tests SKAT-O and KS-Burden.

In addition to the benefits of the KS-Burden test, our approach has also a number of limitations. As shown, given more than 1 causal cluster the KS test loses statistical power. However, the number of causal cluster depends on the sizes of the genomic region as well as the underlying genetic architecture. Furthermore, the use of the combined KS-Burden test is able to, at least partially, recover these shortcomings. Other limitations of the KS-Burden test include its inability to use non-binary phenotypes and make use of covariates as well as variant annotations. While researchers are able to select variants, based on available biological information, and therefore indirect include variant annotations into the test the inability to include covariates is an important limitation. However, most sequencing based studies contain relative homogeneous samples due to potential differences in sequencing platforms and larger population differences in rare variants. Hence despite these limitations the KS-Burden test is a valuable alternative to currently used statistical approaches.

## Conclusion

The KS-Burden test provides better statistical power, compared to most commonly used gene based tests, given a single causal cluster. Furthermore, the test is able to maintain appropriate power in scenarios unfavorable to its underlying assumptions. Hence making it a good alternative to current rare variant tests.

